# Collagen Prolyl Hydroxylases Regulate HIF-α Levels Independently of pVHL in ccRCC

**DOI:** 10.1101/2025.09.14.676157

**Authors:** Yangsook Song Green, Karen Acuña-Pilarte, Marin Jones, Lily Halberg, Thai Huu Ho, Mei Yee Koh

**Author notes:** yCorresponding author Corresponding address: Mei Yee Koh, Department of Pharmacology and Toxicology, University of Utah, 30S 2000E, Salt Lake City, UT 84112.

## Abstract

The hypoxia-inducible factors, HIF-1α and HIF-2α, are master regulators of the hypoxia response. Under ambient conditions, both are hydroxylated by the HIF prolyl hydroxylases (HIF PHDs) resulting in ubiquitination by the pVHL E3 ligase complex, leading to subsequent proteasomal degradation. During hypoxia, the HIF PHDs are inhibited, resulting in HIF-α stabilization and transcriptional activation of genes involved in the adaptation to hypoxia. Previous studies have shown that the collagen PHD, P4HA1, which promotes proline hydroxylation of collagen, inhibits the HIF PHDs by modulating levels of α-ketoglutarate and succinate, thus enhancing HIF-1α stability by preventing pVHL-mediate degradation. Here, we investigate the role of collagen PHDs in the regulation of HIF-1/2α in the setting of pVHL deficiency in ccRCC. We show that the collagen PHDs P4HA1 and P4HA2 are required for HIF-1α translation and HIF-2α transcription and translation independently of pVHL function, through a mechanism regulated in part by P4HA1/2-driven collagen production. Thus, we reveal a novel pVHL-independent mechanism of HIF-1/2α regulation driven by P4HA1/2 in ccRCC. Since the HIFs have been strongly implicated in ccRCC initiation and progression, our data suggests that inhibition of P4HA1/2 may be a promising therapeutic strategy in ccRCC.

## Background

Clear cell renal cell carcinoma (ccRCC) is the most common form of kidney cancer. ccRCC is typically initiated by loss of the von Hippel-Lindau (*VHL*) tumor suppressor gene, resulting in the pseudo-hypoxic activation of the hypoxia inducible factors, HIF-1α and HIF-2α [1-3]. The VHL protein (pVHL) acts as the substrate recognition component of the E3 ligase complex that targets the oxygen labile HIF-1α and HIF-2α subunits for proteasomal degradation under aerobic conditions. The binding of pVHL to HIF-α subunits is contingent upon the iron- and oxygen-dependent hydroxylation of proline residues within HIF-1α and HIF-2α by the HIF prolyl 4-hydroxylases (HIF PHDs) 1-3 [4]. Under hypoxic conditions, iron depleted conditions or in the presence of pVHL mutations, HIF-1α and HIF-2α are stabilized, and enter the nucleus where they heterodimerize with HIF-1β, forming the HIF-1 or HIF-2 transcriptional complexes, respectively. The HIF heterodimers bind to conserved hypoxia response elements within regulatory regions of target genes to activate transcription of hundreds of genes critical for the adaptation to hypoxia, and for tumor progression, such as those promoting aerobic glycolysis, angiogenesis, and metastasis [5, 6]. Constitutive HIF activation, particularly HIF-2α plays a central role in ccRCC progression and the first selective HIF-2α inhibitor, belzutifan, has been FDA approved for the treatment of sporadic ccRCC and cancers associated with VHL disease (a hereditary tumor predisposition syndrome associated with pVHL loss of function) [7] [8, 9]. Although implicated as a tumor suppressor and frequently lost in ccRCC cells, tumor-associated macrophage-specific expression of HIF-1α has been associated with poor prognosis in ccRCC, suggesting that it too may contribute to ccRCC progression [10-12].

Prolyl hydroxylation is a common post-translation modification that modulates protein folding and stability in mammalian cells [13]. Although the stability of the HIF-α subunits in response to oxygen is regulated by cytoplasmic HIF PHDs, the most common hydroxyproline in animal cells is found within collagen, which is regulated by the collagen prolyl 4-hydroxylases (collagen P4Hs) that reside within the endoplasmic reticulum [13]. Collagen P4Hs, like the HIF PHDs, are α_2_β_2_ tetrameric α-ketoglutarate (α-KG)-dependent dioxygenases that catalyzes 4-hydroxylation of proline to promote formation of the collagen triple helix, releasing succinate as a product [14]. Collagen P4Hs have a central role in the biosynthesis of collagens, as 4-hydroxyproline residues are essential for the formation of the collagen triple helix [14]. Both the collagen P4H α(I) and α(II) catalytic subunits (P4HA1 and P4HA2 respectively) associate with the same β-non-catalytic PDI/P4HB subunit to form α(I)_2_β_2_ or α(II)_2_β_2_ tetramers. Intriguingly, collagen P4HA1, by modulating levels of α-KG and succinate, reduces hydroxylation of HIF-1α by the PHDs, enhancing its stability in breast cancer cells [15]. Conversely, P4HA1 and P4HA2 (hereafter P4HA1/2) are activated by HIF-1 and contribute to tumor progression by remodeling the extracellular matrix to promote cancer cell invasion and metastasis [16-18].

However, the impact of P4HA1/2 on the regulation of HIF-1α and HIF-2α in the context of pVHL loss is unclear. Here, we investigate the role of P4HA1/2 in pVHL deficient ccRCC cells and show that P4HA1/2 regulate the HIFs independently of pVHL through a mechanism dependent on P4HA1/2 catalytic activity and/or collagen production.

## Materials and methods

### Immunohistochemistry

Studies were conducted in accordance with the Declaration of Helsinki. Human ccRCC and uninvolved kidney tissue were obtained from archival samples from patients who had provided written informed consent according to protocols approved by the institutional review boards at the Mayo Clinic, AZ. TMAs containing tumor core samples with paired normal from 25 patients with ccRCC pathological grades 2-3 (19 samples with clinical T stage 1 (T1), 1 with T2, and 5 with T3) were stained with antibodies for P4HA1 (66101-1-1g Proteintech, Rosemont IL) and P4HA2 (NBP2-52922, Novus Biologicals, Centennial, CO) at a 1:500 dilution using methods as previously described [12]. Slides were scanned using the Aperio AT2 digital scanning system and cytosolic staining intensity quantitation was performed using Aperio digital imaging software (Leica Biosystems, Buffalo Grove, IL).

### Cell culture and reagents

Renal cell carcinoma cell lines ACHN and 786-0 were purchased from ATCC (Manassas, VA), whereas UOK101 and RCC10 were generous gifts from M. Celeste Simon (University of Pennsylvania). VHL-deficient RCC4 and RCC4 cells with stable re-expression of wild-type VHL were generous gifts from Paul Corn (University of Texas M.D. Anderson Cancer Center). Cells were maintained in Dulbecco’s modified Eagle’s medium (DMEM, ThermoFisher Scientific) supplemented with 10% FBS (Gibco) and maintained in a humidified atmosphere at 37°C in 5% CO2 and routinely tested and verified as mycoplasma negative using the MycoAlert kit (Lonza). Hypoxic incubations were performed at 37 °C in 1% O_2_ using the Whitley H35 Hypoxystation (HypOxygen, Frederick, MO). The selective inhibitor of collagen P4H, diethyl-pythiDC (dp-DC), was purchased from MedChemExpress (Monmouth Junction, NJ), whereas cycloheximide and Rat Tail Collagen were from Sigma-Aldrich (St Louis, MO). After siRNA transfections and treatments, cells were maintained in serum-free media prior to lysis to avoid variability associated with serum on downstream kinase signaling.

### Western blotting

Cells were lysed in ice-cold buffer supplemented with dithiothreitol and protease/phosphatase inhibitors, and total protein concentration was determined using the Pierce BCA Protein Assay Kit (ThermoFisher Scientific, Waltham, MA) according to the manufacturer’s instructions. Samples were prepared at a concentration of 1□mg/ml in sample buffer, resolved by SDS-PAGE, and transferred onto nitrocellulose membranes. Membranes were sectioned based on molecular weight, stripped, and reprobed for proteins within similar molecular weight ranges. Primary polyclonal antibodies used were P4HA1 (NB100-57852) and P4HA2 (NB110-40494) purchased from Novus Biologicals (Centennial, CO), phospho-S6 (4858), S6 (2217), phospho-ERK (4370), ERK (9102), phospho-AKT (4060), AKT (4691), phospho-mTOR (2983), mTOR (2972), AXL (4566), HIF-2a (70862), phospho-tyrosine (8954), GAPDH (5175) and EphA7 (64801) from Cell Signaling Technology (Danvers, MA); phospho-AXL (AF2228) from R&D Systems (Minneapolis, MN); ErbB4 (ab32375) from Abcam (Cambridge, UK); puromycin (MABE343) from MilliporeSigma (Burlington, MA); and HIF-1a (610959) from BD Biosciences (Franklin Lakes NJ). Imaging of the membranes was performed using the Fluorchem M imaging system (Protein Simple, San Jose, CA), and densitometry was performed using multiplex band analysis provided by the AlphaView Software (Protein Simple).

### DNA plasmid and siRNA transfection

Transient transfections were performed 24 hours after seeding at 30-40% confluency. DNA transfections were performed using X-tremeGENE HP (Roche Diagnostics, Indianapolis IN), whereas siRNA was delivered using Lipofectamine™ 2000 (ThermoFisher Scientific, Waltham MA) according to the manufacturer’s protocols. Cells were harvested 48h after DNA transfection, or 72-96 h post siRNA transfection. siRNAs targeting P4HA1 and P4HA2 were ON-Target plus SMARTpool constructs purchased from Dharmacon Discovery (Lafayette, CO) or Ambion™ Silencer™ Select Pre-Designed siRNA (ThermoFisher) or non-targeting controls purchased from each vendor. siRNAs targeting ErbB4, AXL, and EphA7 were ON-Target plus SMARTpool constructs purchased from Dharmacon Discovery.

### Quantitative Reverse Transcription Polymerase Chain Reaction (qRT-PCR)

Total RNA was isolated from RCC4 cells using RNAeasy Mini kit (Qiagen) and quantified using the NanoDrop 2000c (ThermoFisher Scientific). Samples for reverse transcription were prepared using 1 ug of RNA, RT enzyme and buffer mix. Samples for qRT-PCR were prepared using TaqMan™ Universal PCR master mix (Applied Biosystems) and the following probes: *HIF1A* (Hs00153153_m1), *EPAS1* (Hs01026149_m1), or *B2M* (Hs00187842_m1) from ThermoFisher Scientific (Waltham, MA). qRT-PCR experiments were performed on a LightCycler ® 480 (Roche Diagnostics, Indianapolis, IN). Relative mRNA levels were normalized to housekeeping gene *B2M* using the ΔΔCt method.

### Puromycin labelling and Immunoprecipitation

Newly synthesized proteins from cells cultured under serum-free conditions were labeled with puromycin using the SUnSET assay, as previously described [19]. Briefly, 48 hours after siRNA transfection, cells were incubated with 1□µg/mL puromycin (Gibco, A1113803) in complete media (containing FBS) for 30 minutes. Following this labeling period, the medium was replaced with fresh media, and cells were incubated for an additional hour at 37□°C in a humidified incubator with 5% CO_2_. Immunoprecipitation was performed using 1□mg of total protein lysate and the appropriate antibody, followed by overnight incubation with protein A-Sepharose beads (Sigma-Aldrich, P3391) at 4□°C. The following day, samples were washed five times with lysis buffer, then centrifuged, and the resulting bead pellets were resuspended in sample buffer and boiled for 5 minutes at 95□°C, after which samples were analyzed via SDS-PAGE.

### Receptor Tyrosine Kinase Array

Cells were transfected with p4HA1/2 siRNA or control siRNAs, and cultured for 48 hours, then rinsed with PBS and harvested with lysis buffer containing protease inhibitors. Lysates were resuspended, rocked at 4°C for 30 min and centrifuged at 14,000 x g for 15 min. Total protein was quantified using the Pierce BCA Protein Assay Kit (ThermoFisher Scientific) following the manufacturer’s protocol. The Receptor Tyrosine Kinase Array was performed according to the manufacturer’s instructions (ARY001B, R&D Systems, Minneapolis, MN).

### Statistical analysis

Significance between groups was calculated using Students’ T-test using GraphPad Prism 10.4.2 software with p < 0.05 considered significant.

## Results

### P4HA1 and P4HA2 are highly expressed in ccRCC

Since P4HA1/2 are known HIF targets, we examined levels of P4HA1/2 in ccRCC tissue compared to uninvolved normal kidney. Antibody specificity for P4HA1 and P4HA2 was validated using cell pellets of ACHN kidney cancer cells (pVHL wild-type) exposed to normoxia and hypoxia (Supplementary data S1). Having ascertained antibody specificity for P4HA1 and P4H2, we stained a tumor microarray containing paired ccRCC and paired uninvolved kidney tissue for P4HA1 and P4HA2. We found that ccRCC tumors showed significantly higher levels of P4HA1/2 protein compared to the adjacent uninvolved kidney (**Fig. 1A** and **1B**), as expected. Both P4HA1 and P4HA2 levels were diffusely expressed in the cytoplasm of ccRCC tissue but were largely negative within uninvolved kidney. When we examined levels of P4HA1/2 in ccRCC cells lines, we found that VHL-deficient ccRCC cell lines expressed much higher levels of P4HA1 and P4HA2 in normoxia compared to a VHL wild type ACHN cells, possibly due to constitutive HIF-1/2α expression associated with pVHL loss (**Fig. 1C**). However, when the cells were exposed to increasing durations of hypoxia, we found a general increase in P4HA1/2 protein levels with increasing hypoxic duration in both pVHL wild-type and pVHL deficient cells (with the exception of RCC4 cells; **Fig. 1C**). Intriguingly, P4HA1/2 levels were maximally stabilized under prolonged hypoxia (48h), which was past the peak of HIF-1α induction of any of the cell lines examined (**Fig. 1C)**, suggesting that P4HA1/2 are regulated by HIF-α-dependent and independent mechanisms, including through a translational mechanism which has previously been described [20].

**Figure 1.**
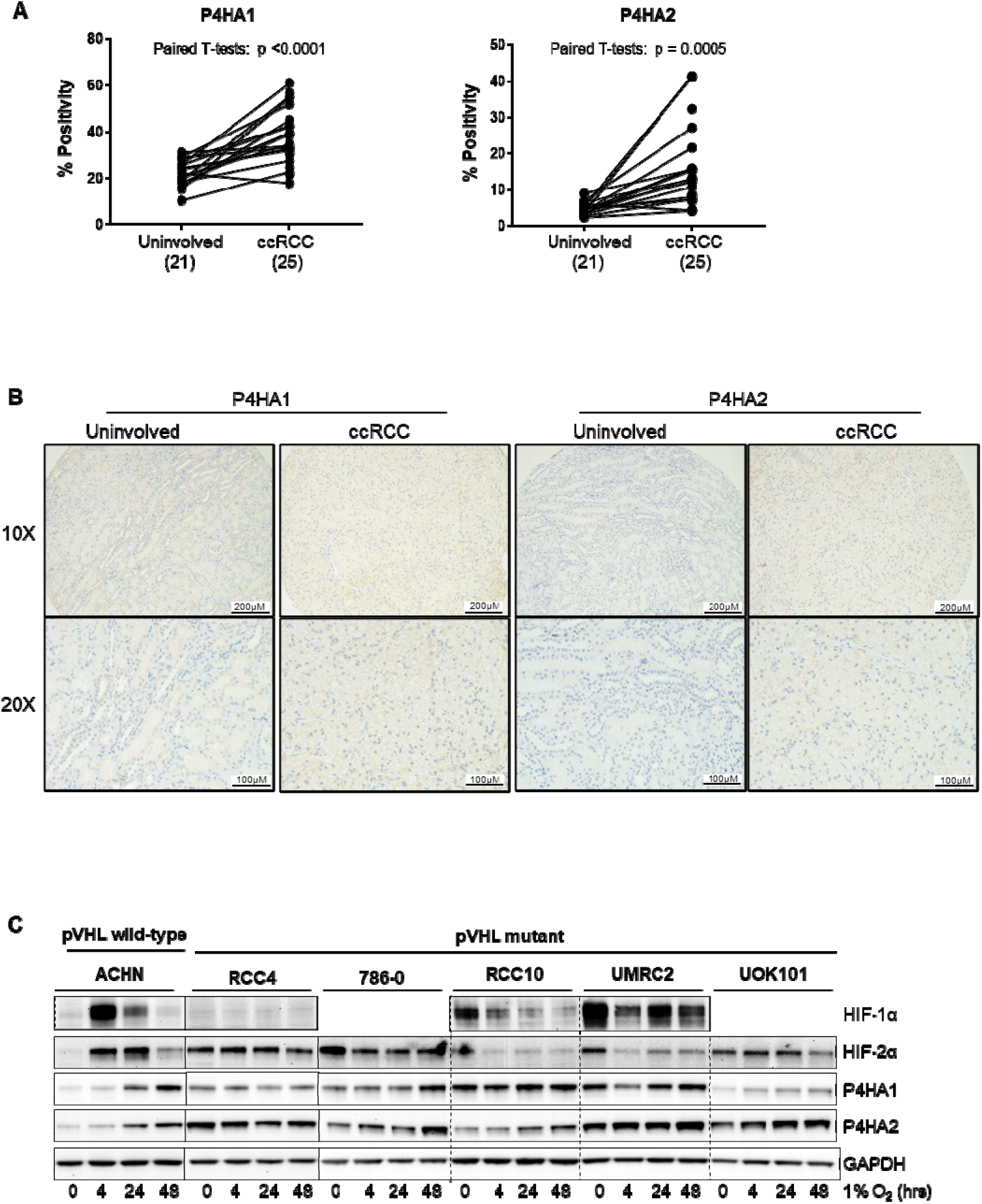
P4HA1 and P4HA2 are highly expressed in ccRCC. A) IHC quantification of P4HA1 and P4HA2 staining in ccRCC tumors and paired adjacent uninvolved normal kidney tissue. Number of cases analyzed are indicated in brackets. Each data point represents an individual case. B) Representative images of P4HA1 and P4HA2 in uninvolved and ccRCC tissue at 10X and 20X V magnification. C) Western blot analysis showing the time-dependent effects of hypoxia on HIF-1α, HIF-2α, P4HA1, and P4HA2 expression in a panel of ccRCC cell lines with either wild-type or mutant p HL. Solid lines indicate demarcation of blots from separate gels. Dotted lines are provided for clarity to indicate bands from the same gel.

### P4HA1 and P4HA2 regulate HIF-1α and HIF-2α in a pVHL independent manner

Since P4HA1 was previously shown to induce HIF-1α by decreasing prolyl hydroxylation, thus blocking subsequent pVHL-dependent degradation, we investigated whether knockdown of P4HA1/2 or the non-catalytic unit of the P4H tetramer, P4HB, could impact HIF-α levels in pVHL deficient cells [15]. Consistent with previous reports in breast cancer, we found that transfection with P4HA1/2 siRNA decreased HIF-1α levels, and to differing extents, also HIF-2α levels in pVHL wild-type cells in hypoxia (ACHN and RCC4 VHL cells; **Fig 2A-B**) [15]. Intriguingly, we found that P4HA1/2 siRNA also markedly reduced levels of HIF-2α and HIF-1α to differing extents in a panel of pVHL deficient cells using siRNAs for P4HA1/2 purchased from different suppliers (Ambion and Dharmacon; **Fig. 2B-C;** hereafter siRNAs from Dharmacon were used). The decreases in HIF-α in both pVHL wild-type and pVHL mutant cells were associated with decreases in phosphorylation of ribosomal protein S6 (p-S6, S235/236) and was also observed when P4HB was knocked down (**Fig. 2A-C**). We observed similar decreases in levels of p-ERK with P4HA1/2 siRNA, whereas levels of p-AKT (T308, S473) and p-mTOR (S2448) were unchanged (**S2A-C**). We also observed decreases in HIF-1/2α levels when cells were treated with the selective inhibitor of collagen P4Hs, diethyl-pythiDC (dp-DC) that inhibits collagen P4H activity at concentrations that do not cause iron deficiency (**Fig. 2D**) [21]. Here, treatment with dp-DC decreased levels of HIF-1/2α, p-S6 and p-ERK (although changes in HIF-1α and p-ERK were more subtle) but did not alter levels of p-AKT and p-mTOR, suggesting that P4H activity promotes HIF-α synthesis and decreases S6 and ERK phosphorylation (**Fig. 2D, S2D**). However, overexpression of P4HA1/2, while increasing p-S6, did not change levels of HIF-1/2α in pVHL deficient RCC4 cells suggesting that HIF-1/2α may already be maximal in this pVHL-deficient cell type (**Fig. 2E**). By contrast, P4HA1/2 overexpression modestly increased levels of HIF-1α in pVHL wild-type ACHN cells without changing levels of p-S6 (**Fig. 2F**) in agreement with previous reports [15]. Consistent with the role of P4Hs in the production of collagen, we found that the addition of exogenous rat tail collagen attenuated the decreases in HIF-1/2α and in p-S6 mediated by P4HA1/2 siRNA especially in normoxia, suggesting these effects were due at least in part, to the decreased collagen production associated with P4HA1/2 siRNA (**Fig. 2G**). Taken together, our results suggest that P4HA1/2 are necessary for the maintenance of HIF-2α and to a lesser extent, HIF-1α, in pVHL-deficient ccRCC cell lines through a mechanism which is associated with S6 and ERK phosphorylation but independent of mTOR.

**Figure 2.**
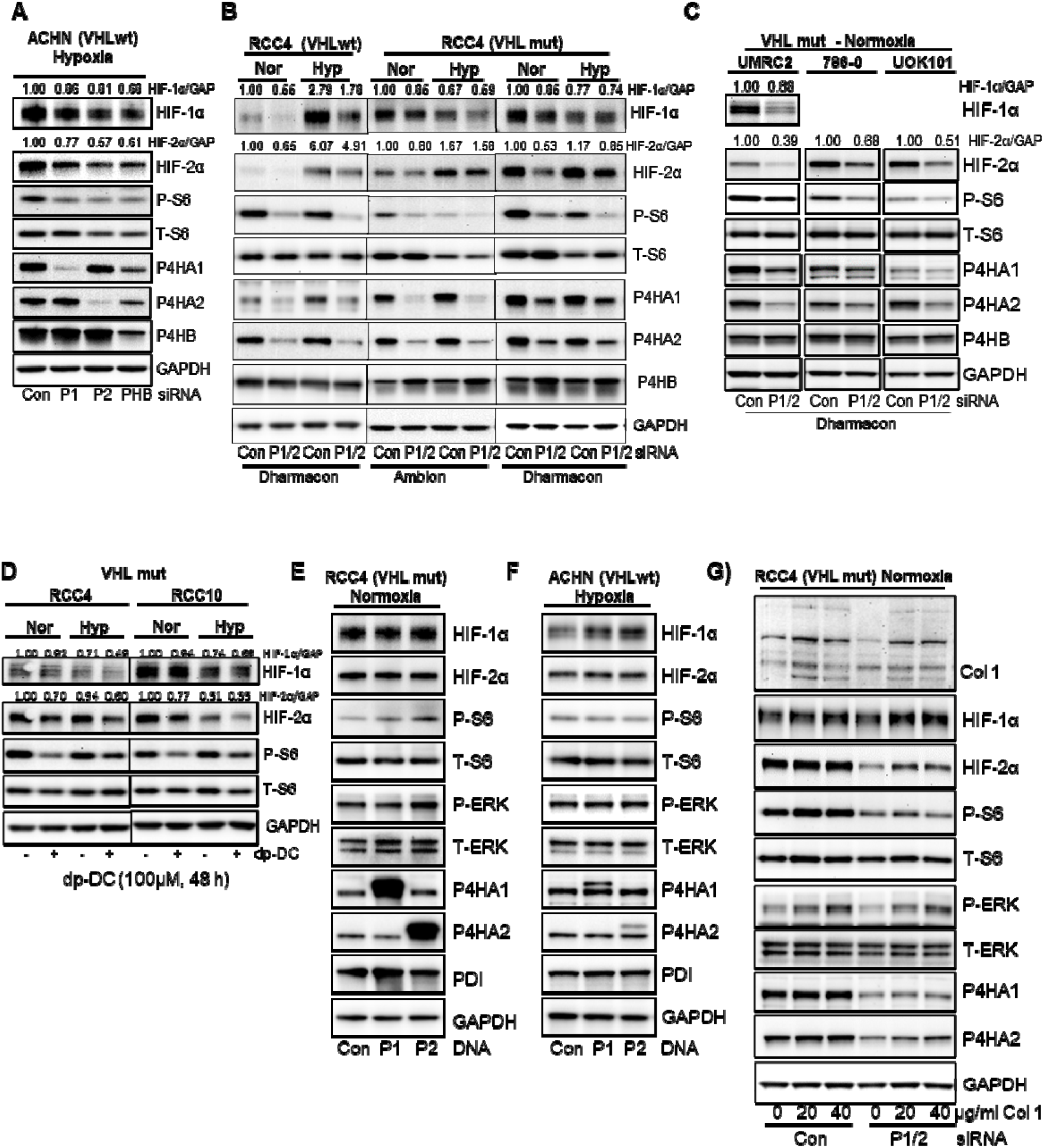
P4HA1 and P4HA2 are required to maintain HIF-1α and HIF-2α protein levels in normoxia and hypoxia independently of pVHL. A) Western blot analysis showing the effect of knockdown of P4HA1 (P1), P4HA2 (P2), and P4HB (P4B) or non-targeting control (Con) siRNA on HIF-1α, HIF-2α, and phosphorylated S6 (S235/236) (P-S6) in ACHN cells (VHL wild-type, VHLwt) in normoxia or hypoxia for the last 16 hours prior to lysis. B) Western blot analysis showing the combined effect of knockdown of P4HA1 and P4HA2 (P1/2) using siRNAs from Dharmacon and Ambion, on HIF-1α, HIF-2α, and P-S6 in RCC4 cells expressing mutant (VHL mut) or wild-type pVHL in normoxia or hypoxia for the last 24 hours prior to lysis. C) Western blots showing the combined effect of P4HA1 and P4HA2 knockdown on HIF-1α, HIF-2α, and P-S6 in three VHL-mutant ccRCC cell lines, UMRC2, 786-0, and UOK101 in normoxia. D) Western blot analysis of RCC4 and RCC10 cells treated with the collagen prolyl-4-hydroxylase inhibitor diethyl-pythiDC (dp-DC) for 48 hours under normoxic or hypoxic (4hrs) conditions. E, F) Western blot analysis of RCC4 cells (pVHL mutant) under normoxic conditions, (E), and ACHN cells (pVHL wild-type) exposed to hypoxia for 20 hours (F), showing the effects of P4HA1 (P1) and P4HA2 (P2) overexpression on P-S6, P-ERK, HIF-1α, and HIF-2α levels. G) Western blot analysis of RCC4 cells (pVHL mutant) treated with collagen 1 (Col 1) for 24 hours under normoxic conditions after transfection with control (Con) siRNA or siRNA targeting P4HA1 and P4HA2 (P1/2). Solid lines indicate demarcation of blots from separate gels.

### P4HA1/2 regulate HIF-1/2α at the level of transcription and translation

To determine the mechanism by which P4HA1/2 knockdown decreases HIF-1/2α levels, we investigated the effects of P4HA1/2 siRNA on the transcription and translation of HIF-1/2α. Intriguingly, we found that P4HA1/2 knockdown significantly decreased HIF-2α (*EPAS1*) transcription in both RCC4 and 786-0 cells (**Fig. 3A-B**). By contrast, HIF-1α transcription was significantly increased in RCC4 cells with P4HA1/2 siRNA (**Fig. 3A**). The impact of P4HA1/2 siRNA on HIF-1α transcription could not be assessed in 786-0 cells since they do not express HIF-1α transcript or protein. Since our data also showed that P4HA1/2 siRNA decreased the phosphorylation of S6, a component of the 40S ribosomal subunit known to be involved in translation, we investigated the impact of P4HA1/2 siRNA on HIF-1/2α at the level of translation using the SUnSET assay which measures puromycin incorporation as an indicator of translation [19]. Cells transfected with P4HA1/2 siRNA for 48 hours (a time point prior to maximal loss of HIF-α protein to enable sufficient HIF-α for subsequent immunoprecipitation) were pulsed with puromycin, then lysed after which HIF-1α or HIF-2α was immunoprecipitated and probed for the incorporation of puromycin via western blotting. Indeed, we found that P4HA1/2 knockdown was associated with markedly decreased puromycin incorporation into both HIF-1α and HIF-2α (**Fig. 3C-E**), suggesting the P4HA1/2 is required for the translation of both HIF-1α and HIF-2α. Taken together, the data suggests that P4HA1/2 knockdown decreases HIF-1α translation, and inhibits both the transcription and translation of HIF-2α. This may explain the more pronounced effect of P4HA1/2 knockdown in decreasing HIF-2α compared to HIF-1α.

**Figure 3.**
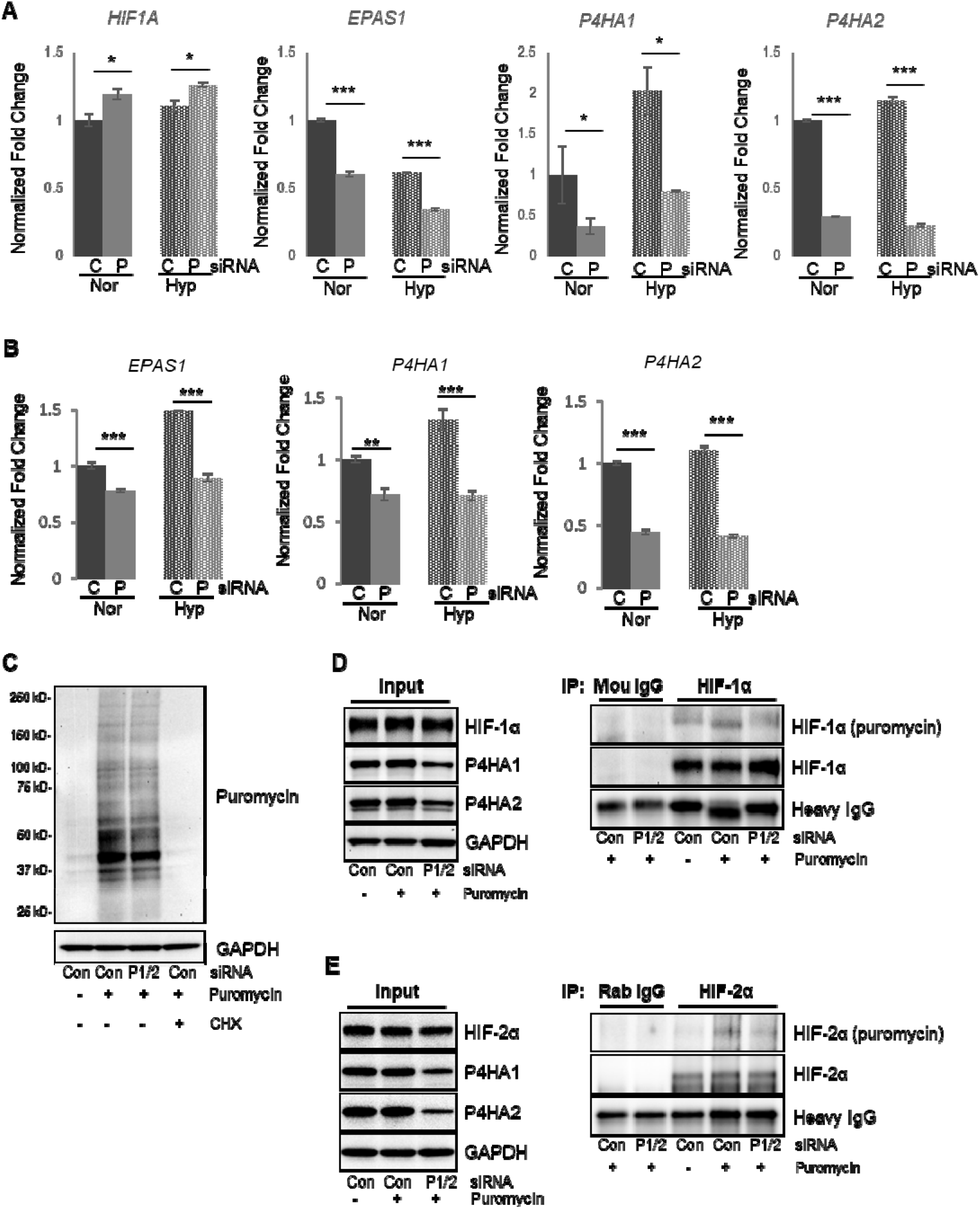
P4HA1 and P4HA2 knockdown decreases HIF-1α translation and HIF-2α transcription/translation. A, B) Quantitative qRT-PCR analysis of *HIF1A, EPAS1* (encoding HIF-2α), *P4HA1*, and *P4HA2* mRNA levels in RCC4 (A) and 786-0 (B) cells under normoxic and hypoxic (24 hours) conditions in serum-free media 96 hours after transfection with non-targeting control (C) siRNA or siRNA for P4HA1 and P4HA2 (P). C) Western blot analysis of puromycin-labeled proteins following combined knockdown of P4HA1 and P4HA2 (P1/2) or control (Con) in RCC4 cells, treated with puromycin and cycloheximide (CHX) under normoxic conditions. D, E) Western blot analysis of input and immunoprecipitated samples using HIF-1α (D) or HIF-2α (E) antibodies and then probed with puromycin antibody in RCC4 cells with combined knockdown of P4HA1 and P4HA2, treated with puromycin shown in C.

### P4HA1/2 regulation of HIF-1α and HIF-2α does not depend on receptor tyrosine Kinases, AXL and EphA7

Since components of the extracellular matrix including collagen play important roles in activating cell surface receptors to initiate signaling pathways, we utilized a receptor tyrosine kinase (RTK) array to identify potential RTKs perturbed by P4HA1/2 knockdown in RCC4 cells [22]. We detected only six RTKs that showed detectable phosphorylation levels in this assay, of which, three RTKs, ErbB4, AXL and EphA7, showed reduced phosphorylation after knockdown of P4HA1/2 in RCC4 cells (**Fig. 4A-B**). Of note, there was no impact of P4HA1/2 knockdown on overall levels of phospho-tyrosine containing proteins as detected by a pan-phospho-tyrosine antibody (**Fig. 4C**). We thus investigated the potential involvement of ErB4, Axl and EphA7 in mediating the effects of siP4HA1/2 on HIF1-2α levels. We could not detect a band corresponding to full length ErbB4 in RCC4 cells (despite observing a band of the expected size in HEK293 cells), which is consistent with a previous report indicating that ErbB4 expression is downregulated in ccRCC (**S3A**) [23]. These suggest that the ErbB4 detected in the RTK array may be non-specific. By contrast, we found that EphA7 was only detectable in RCC4 cells (**S3B**), but not in other ccRCC cell lines RCC10 and 786-0, whereas AXL was detected across all three ccRCC cell lines. To investigate the effects of P4HA1/2 on the phosphorylation of AXL and EphA7, we performed immunoprecipitations of AXL and EphA7 in RCC4 cells transfected with non-targeting (control) or P4HA1/2 siRNA and probed these with an antibody for pan-phospho-tyrosine. Indeed, we found that P4HA1/2 siRNA markedly decreased the phosphorylation of AXL and EphA7 (**Fig. 4D-E**). However, siRNA knockdown of AXL or EphA7 did not independently decrease levels of HIF-1α, HIF-2α, P-S6 and pERK suggesting that the effects of P4HA1/2 knockdown on these proteins may not be solely driven by these proteins but may be mediated by the downregulation of multiple different RTKs and potentially other mediators (**Fig 4F-G**). Taken together, our findings reveal a novel regulatory pathway of HIF-1/2α mediated by collagen P4HA1/2 potentially mediated at least in part, by activation of RTKs.

**Figure 4.**
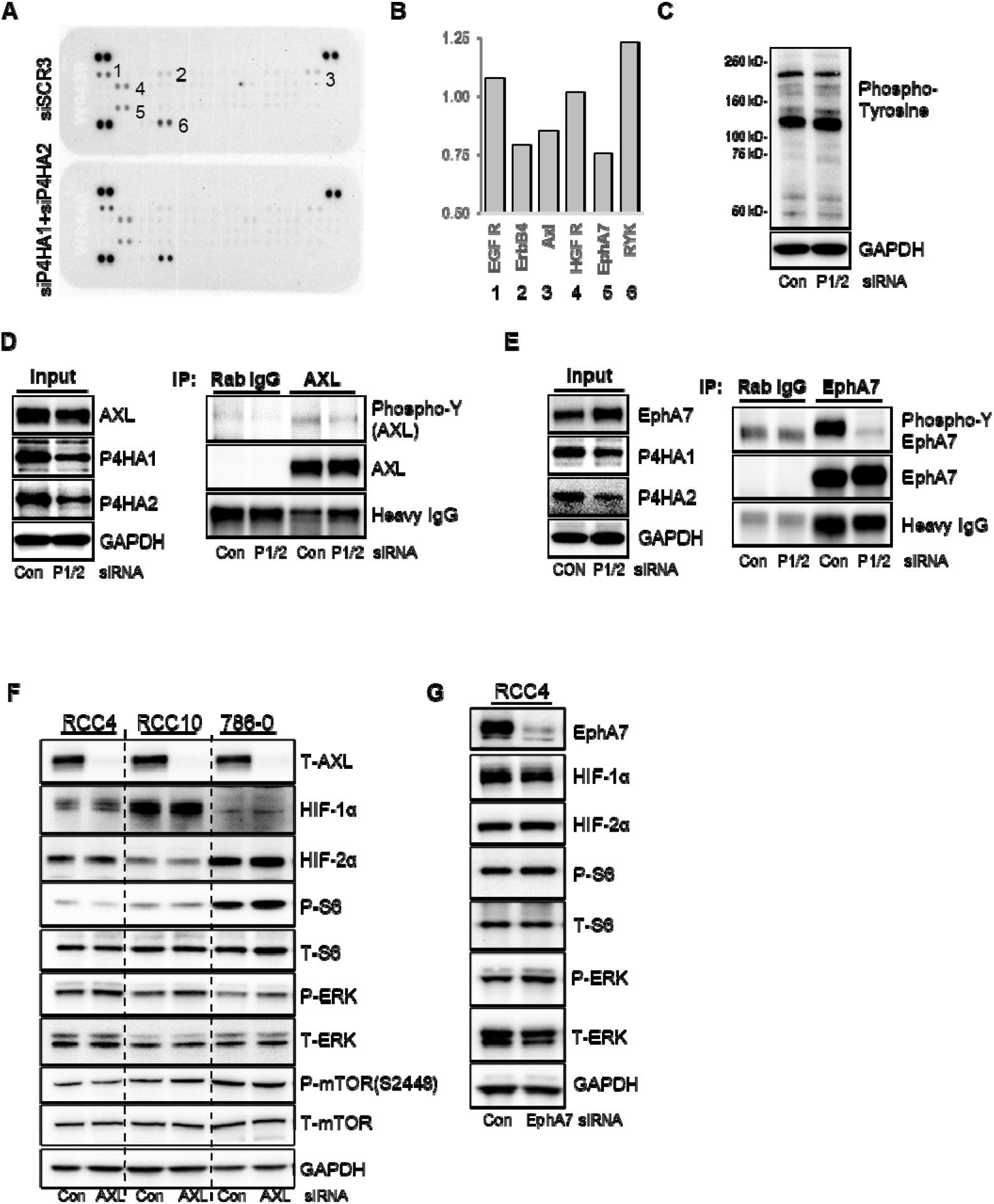
P4HA1 and P4HA2 knockdown decrease phosphorylation of receptor tyrosine kinases (RTKs). A) RTK array showing the phospho-tyrosine levels of EGF receptor (1), ErbB4 (2), AXL (3), HGF receptor (4), EphA7 (5), and RYK (6) 48 hours after combined knockdown of P4HA1 and P4HA2 (P1/2 siRNAs) or control (Con) siRNA in RCC4 cells in normoxia. B) Quantification of the RTK array in (A). C) Western blot analysis of overall tyrosine phosphorylation levels in RCC4 cells following combined knockdown of P4HA1 and P4HA2. D, E) Western blot analysis of input and immunoprecipitated samples using AXL (D) or EphA7 (E) antibodies and then probed with phospho-Tyrosine (phospho-Y) antibody in RCC4 cells with combined knockdown of P4HA1 and P4HA2 under normoxic conditions. F) Western blot analysis of the impact of AXL knockdown (72 hours) on protein levels of HIF-1α, HIF-2α, and phosphorylation of S6, and ERK in RCC4, RCC10, and 786-0 cells in normoxia. G) Western blot showing the effect of EphA7 knockdown (72 hours) on protein levels of HIF-1α, HIF-2α, and phosphorylation of S6 and ERK in RCC4 cells in normoxia. Dotted lines are provided for clarity to indicate bands from the same gel.

## Discussion

ccRCC is uniquely initiated by loss of function mutations in the *VHL* gene, which results in the constitutive activation of HIF-1α and HIF-2α. HIF-1α and HIF-2α activate both distinct and overlapping genes critical for response to hypoxia including those involved in angiogenesis as well as metabolic and immune reprogramming that can contribute to tumor progression [24]. Here we show that the HIF target genes, P4HA1/2, regulate both HIF-1α and HIF-2α independently of pVHL through a mechanism dependent on P4HA1/2 catalytic activity and/or collagen production by regulating their translation (HIF-1α and HIF-2α) and transcription (HIF-2α only). This likely mediates a feed-forward loop of HIF and P4HA1/2 activation that may contribute to further increases in HIF and P4HA1/2 levels.

In breast cancer cells, P4HA1-mediated hydroxylation of collagen decreases the availability of α- ketoglutarate, which is required for HIF PHD-mediated hydroxylation of HIF-1α. This reduces HIF-1α hydroxylation and subsequent pVHL-mediated degradation, stabilizing HIF-1α protein levels [15]. P4HA2 has also been shown to stabilize HIF-1α through a similar mechanism in bladder cancer, whereas either P4HA1 or P4HA2 knockdown decreases HIF-1α in pancreatic cancer cells [18, 25]. Although these studies have not specifically addressed the impact of P4HA1/2 on HIF-2α in these settings, since HIF-2α is similarly regulated by the HIF PHDs, it is likely that P4HA1/2 similarly contribute to the stability of HIF-2α. Although these previous studies have linked P4HA1/2 to HIF-1α (and potentially HIF-2α) through P4HA1/2’s competition with the HIF PHDs, here we show for the first time that P4HA1/2 regulates the HIFs independently of pVHL (and thus the HIF PHDs) by inducing transcription and translation through a mechanism dependent on P4HA1/2 catalytic activity and/or collagen production. We show that the effects of P4HA1/2 on HIF-α transcription and/or translation are associated with ERK1/2 and ribosomal protein S6 activation. ERK1/2, a member of the MAPK family, plays a central role in transmitting extracellular signals through phosphorylation-driven signaling cascades [26]. Multiple agonists such as growth factors, cytokines and extracellular matrix proteins such as collagen, can activate relevant receptors such as EGFR or VEGFR, resulting in activation of downstream kinases such as Ras, which activates a signaling cascade leading to ERK phosphorylation, which in turn activates a plethora of downstream signaling pathways related to cell proliferation, differentiation and survival. ERK phosphorylation has been shown to increase phosphorylation of ribosomal protein S6, a major regulator of protein synthesis, which has been linked to increased HIF-1α translation [27]. Independent of S6, ERK activation by reactive oxygen species has been shown to induce HIF-1α transcription in cancer cells [28]. Although there is no evidence for direct effects of ERK and S6 on HIF-2α expression, our overall findings suggest that activated ERK and S6 associated with P4HA1/2 activity/collagen production may contribute to the transcription and/or translation of HIF-1α and HIF-2α in ccRCC.

Collagen is the most abundant protein in the extracellular matrix (ECM), a network of extracellular macromolecules that plays an important role in maintaining tissue structure and cellular function. Apart from its known structural role in the ECM, collagen also functions as a signaling molecule by activating receptors including Integrin receptors and Discoidin Domain Receptors (DDR1/2), which result in activation of downstream signaling cascades including ERK and S6 [29-33]. The fact that the reduced p-ERK and p-S6 levels associated with P4HA1/2 siRNA were partially rescued by the addition of collagen 1 into cell culture media suggest that the effects of P4HA1/2 siRNA in decreasing HIF-α levels may be driven in part, through impaired collagen-induced signaling (**Fig. 2G**). Although we were unsuccessful in identifying an individual RTK responsible for driving the reduced HIF-α levels associated with siRNA to P4HA1/2, our data suggest that P4HA1/2 may promote the activation of RTKs (and other unknown effectors), in part through increased collagen production, to contribute to the synthesis of HIF-1/2α (**Fig. 5**), at least in the *in vitro* setting. Further studies are required to determine whether they similarly regulate the HIFs in a biologically significant manner in the setting of the tumor microenvironment.

**Figure 5.**
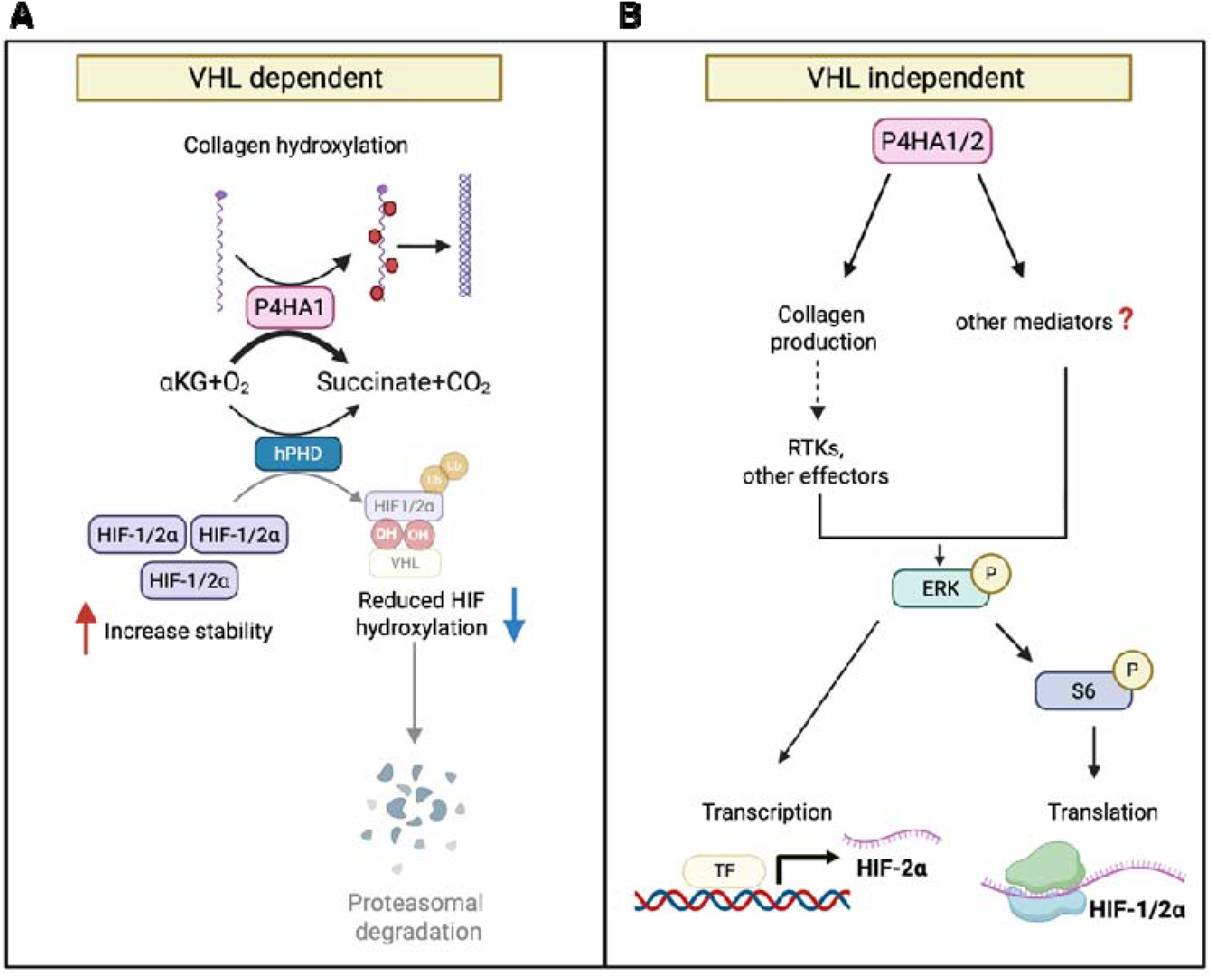
P4HA1/2 promote HIF-1/2α synthesis through pVHL-dependent and PVHL-independent mechanisms. A) P4HA1/2 enhances HIF-1/2α stability by reducing the availability of α-ketoglutarate (αKG), a cofactor required for HIF PHD (hPHD) activity. Reduced αKG availability reduces hPHD-mediated hydroxylation of HIF-1/2α, preventing pVHL-mediated ubiquitination and proteasomal degradation of HIF-1/2α. B) PAHA1/2 also promotes phosphorylation of ERK and S6, potentially through increased collagen production, which stimulates signaling though receptor tyrosine kinases (RTKs) and other mediators. This leads to increased transcription of HIF-2α and enhanced translation of both HIF-1α and HIF-2α that result in higher HIF-1/2α levels independently of pVHL function.

A variety of collagen family member genes are upregulated in ccRCC and high expression levels of COL1A1, COL5A1 and COL6A3 were significantly correlated with decreased overall survival in the TCGA KIRC database [34]. Although deregulation of collagen family members has been associated with increasing metastasis and immune evasion, our data suggest that collagen production mediated by P4HA1 and P4HA2 may also contribute to the maintenance of HIF-1α and HIF-2α, and thus contribute to the progression of ccRCC, and potentially of other tumor types. Indeed, it is possible that the reported efficacy of P4HA1/2 inhibition in mouse tumor models could be at least partly attributed to decreased HIF-1/2α production [35]. Thus, P4HA1/2 inhibition may be a promising therapeutic strategy to both inhibit the pro-invasive/metastatic ECM and decrease HIF-1/2α in both pVHL deficient and pVHL wild-type cells. Our findings also provide a rationale for the combination of P4HA inhibition with belzutifan-mediated HIF-2α inhibition in ccRCC to circumvent resistance associated with gatekeeper mutations in HIF-2α [36].

## Author contributions

YSG performed the experiments, prepared the figures and contributed to the initial draft of the paper; KAP prepared the figures, contributed to data analysis and wrote the manuscript; MJ and LH performed experiments and contributed to data analysis, THH provided human tissue, contributed to data analysis, and provided input on the manuscript; MYK conceived of the project, designed experiments, contributed to data analysis and wrote and finalized the manuscript. All authors reviewed the manuscript.

## Acknowledgements and funding sources

Supported by NIH NCI grants R01CA181106 and DoD KC230225 to MYK and by NIH NCI grants R01CA271503 and NIH-R01CA224917 to THH. Supported in part by funding from Hollings Cancer Center’s Cancer Center Support Grant P30 CA138313 at the Medical University of South Carolina. The content is solely the responsibility of the authors and does not necessarily represent the official views of the NIH

