## Supplemental Figures for "Collagen Prolyl Hydroxylases Regulate HIF-α Levels Independently of pVHL in ccRCC"

### Slide 1
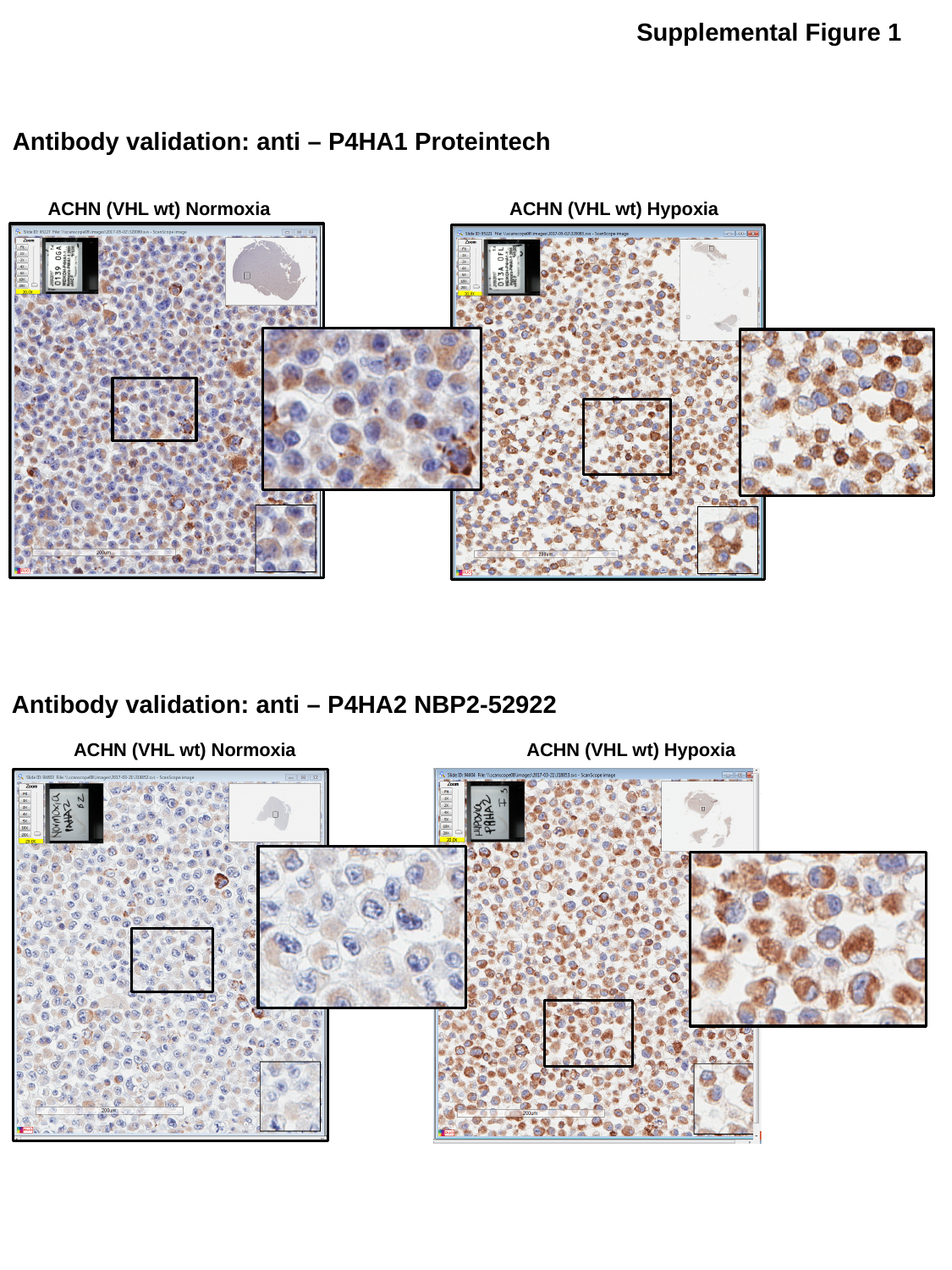

Supplemental Figure 1
Antibody validation: anti – P4HA1 Proteintech
ACHN (VHL wt) Normoxia
ACHN (VHL wt) Hypoxia
Antibody validation: anti – P4HA2 NBP2-52922
ACHN (VHL wt) Normoxia
ACHN (VHL wt) Hypoxia

### Slide 2
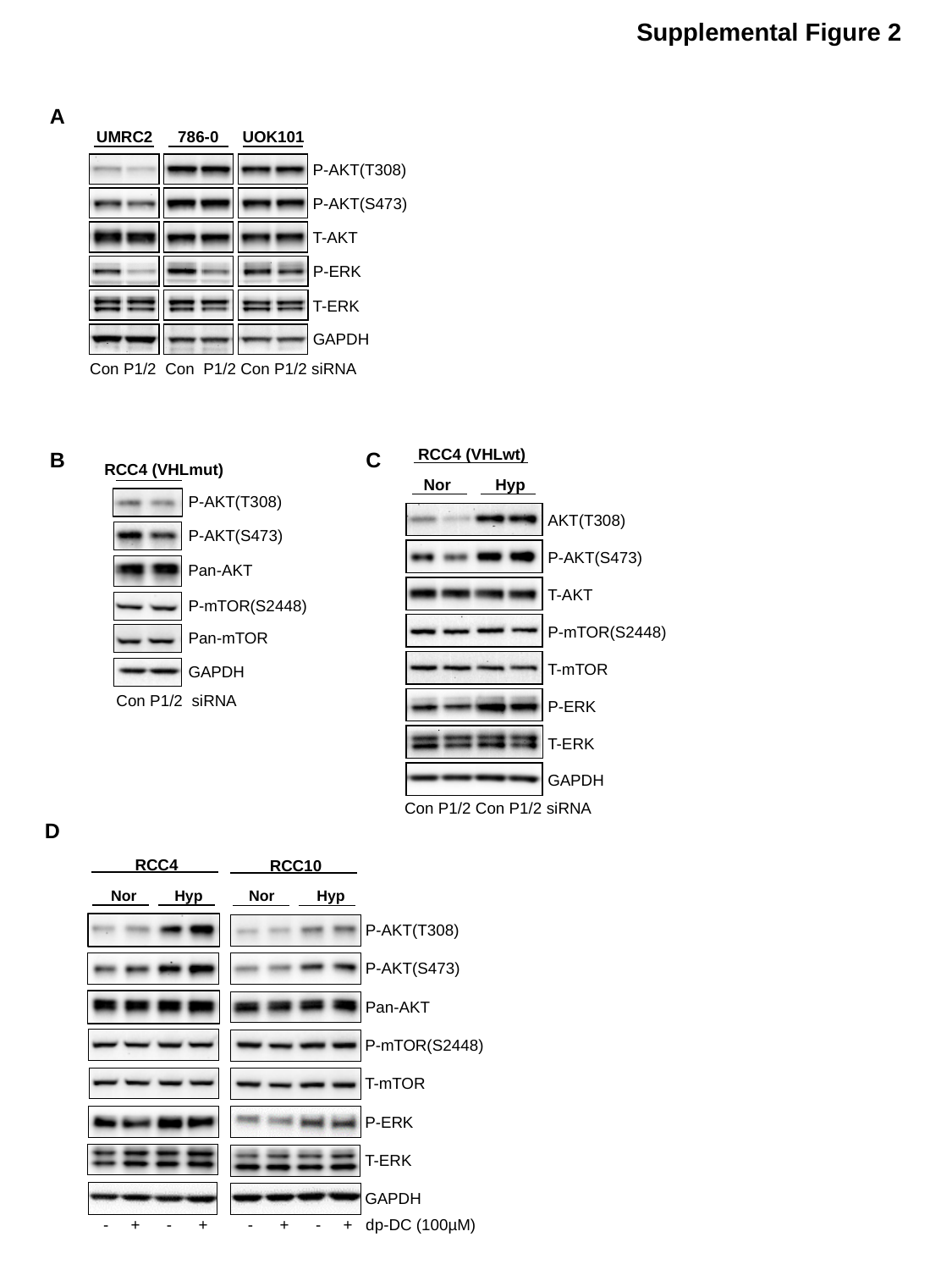

Supplemental Figure 2
A
UMRC2
786-0
UOK101
P-AKT(T308)
P-AKT(S473)
T-AKT
P-ERK
T-ERK
GAPDH
Con P1/2 Con P1/2 Con P1/2 siRNA
RCC4 (VHLwt)
Nor Hyp
AKT(T308)
P-AKT(S473)
T-AKT
P-mTOR(S2448)
T-mTOR
P-ERK
T-ERK
GAPDH
Con P1/2 Con P1/2 siRNA
B C
RCC4 (VHLmut)
P-AKT(T308)
P-AKT(S473)
Pan-AKT
P-mTOR(S2448)
Pan-mTOR
GAPDH
Con P1/2 siRNA
D
RCC4
RCC10
Nor Hyp
Nor Hyp
P-AKT(T308)
P-AKT(S473)
Pan-AKT
P-mTOR(S2448)
T-mTOR
P-ERK
T-ERK
GAPDH
 - + - + - + - + dp-DC (100µM)

### Slide 3
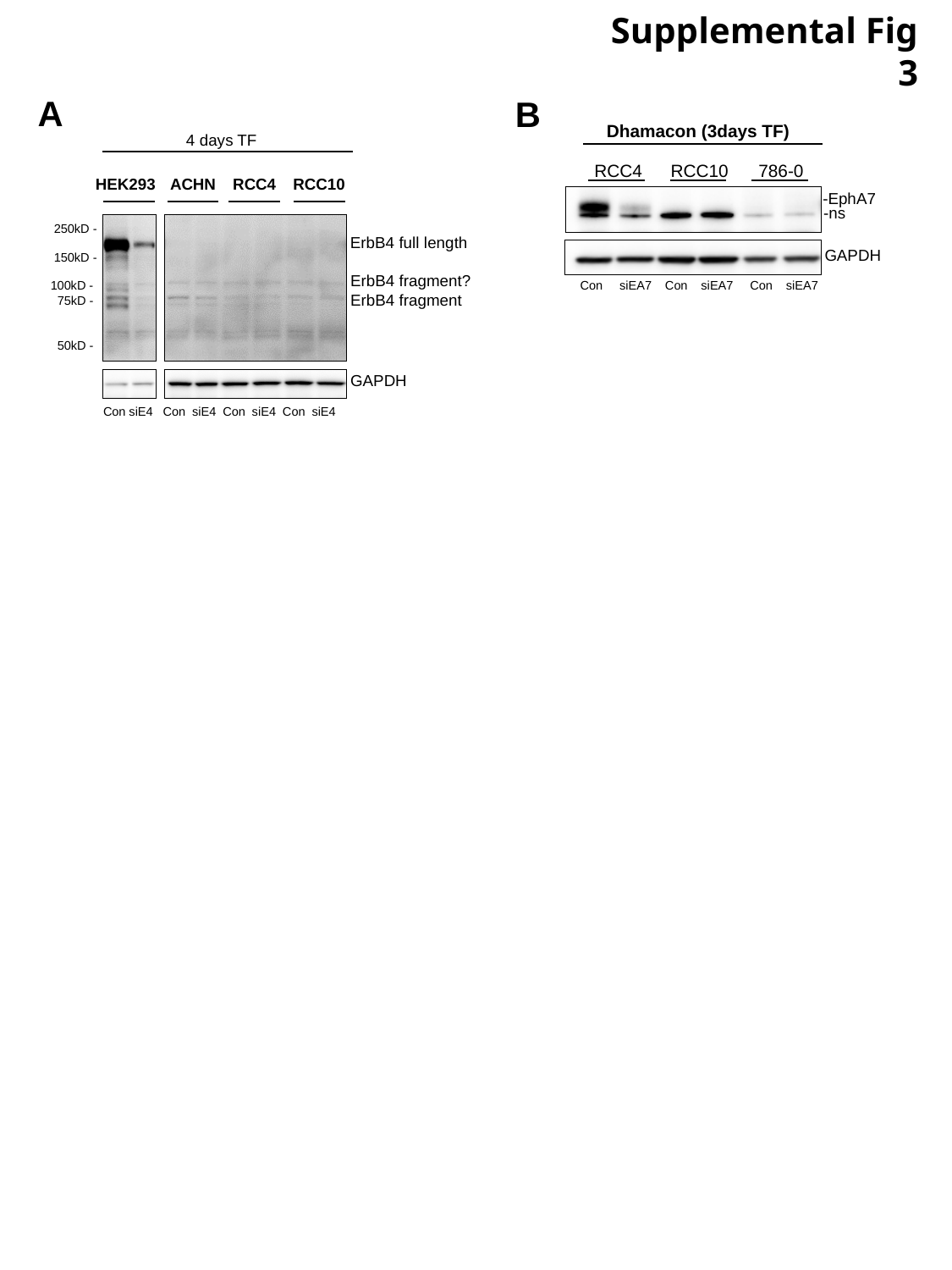

Supplemental Fig 3
A
B
Dhamacon (3days TF)
RCC4
RCC10
786-0
4 days TF
HEK293
ACHN
RCC4
RCC10
250kD -
150kD -
100kD -
75kD -
50kD -
ErbB4 full length
ErbB4 fragment?
ErbB4 fragment
GAPDH
Con siE4 Con siE4 Con siE4 Con siE4
-EphA7
-ns
GAPDH
Con siEA7 Con siEA7 Con siEA7
