## Supplemental figure legends for "Collagen Prolyl Hydroxylases Regulate HIF-α Levels Independently of pVHL in ccRCC"

**Supplemental Figure 1: Validation of antibodies used for immunohistochemistry staining of P4HA1 and P4HA2 in tumor microarrays.** Staining conditions for each antibody was optimized using staining controls of formalin-fixed paraffin embedded normoxic and hypoxic ACHN (VHl wt) cell pellets. Representative images show staining patterns of P4HA1 and P4HA2 in these pellets under the staining conditions used.

**Supplemental Figure 2: Impact of P4HA1/2 siRNA on AKT, mTOR and ERK activation.** A) Western blot analysis of the combined effect of P4HA1 and P4HA2 siRNA on levels of phosphorylated ERK and AKT in UMRC2, 786-0, and UOK101 cells in normoxia. B) Impact of P4HA1 and P4HA2 knockdown on AKT and mTOR phosphorylation in RCC4 cells in normoxia. C) Western blot analysis of the effect of P4HA1 and P4HA2 knockdown on phosphorylation of AKT, mTOR, and ERK under normoxic and hypoxic (24 hourrs) conditions in RCC4 VHL cells. D) Western blot analysis showing the effect of the collagen P4H inhibitor diethyl-pythiDC (dp-DC) on phosphorylation levels of ERK and AKT in VHL mutant RCC4 and RCC10 cells under normoxic and hypoxic (4 hours) conditions.

**Supplemental Figure 3. Expression of ErbB4 and EphA7 in ccRCC cells.** A) Western blot analysis of ErbB4 protein levels following ErbB4 siRNA transfection (siE4) in HEK293, ACHN, RCC4, and RCC10 cells. B) Western blots showing protein levels of EphA7 after EphA7 siRNA transfection (siEA7) in RCC4, RCC10, and 786-0 cells.
